# Skin-Related Effects of Corynebacterium amycolatum SB-1 Ferment Filtrate and Functional Characterization of Ellagic Acid

**DOI:** 10.64898/2026.09.02.748710

**Authors:** Songmi Kim, Seyoung Mun, Misun Kim, Min-Ji Kim, Dong-Geol Lee, Seunghyun Kang, HyungWoo Jo, Kyudong Han

## Abstract

Previous research has established a link between the genus *Corynebacterium* in the skin microbiota and skin health; however, studies investigating the specific effects of *Corynebacterium*-derived substances on skin cells remain limited. In this study, we isolated eight strains of *Corynebacterium* from the facial skin of female participant. To investigate the potential skin improvement effects of these isolates, three cultured skin cell lines (Hs68, HaCaT, and B16-F10) were treated with the supernatants of each strain. We then evaluated and compared the gene expression levels (*ELN*, *COL1A1*, *IL-1β*, and *HAS3*) alongside their inhibitory effects on melanin biosynthesis. *Corynebacterium amycolatum* SB-1 was selected as the final candidate for subsequent analysis, primarily based on its rapid growth rate and high cell yield. Using LC-TOF/MS, we screened the postbiotic candidates from the SB-1 strain and identified ellagic acid as a known polyphenolic antioxidant with potent anti-melanogenic effects beneficial for skin improvement. To validate the effects of ellagic acid on skin cells, we assessed *FLG* gene expression and its inhibition of melanin biosynthesis. Additionally, RNA sequencing was performed to elucidate the underlying molecular mechanisms, revealing that ellagic acid is significantly involved in inhibiting melanogenesis and promoting collagen synthesis.

Taken together, our results highlight ellagic acid derived from strain SB-1 as a highly effective bioactive compound for improving skin health, specifically targeting hyperpigmentation and skin aging.

## Introduction

The cosmetics industry has traditionally focused on developing products that address skin health and aesthetics through synthetic ingredients, emollients, and active compounds. However, recent advancements in microbiome research have shifted the industry’s perspective toward leveraging the skin’s natural microbial ecosystem to enhance skin health and beauty (1). The skin microbiome—comprising bacteria, fungi, viruses, and other microorganisms— plays an integral role in maintaining skin barrier function, modulating immune responses, and protecting against pathogenic invasions. Increasingly, research indicates that the skin’s microbiome may be crucial for developing next-generation cosmetic products that align with the skin’s natural biology (1, 2).

In recent years, the potential for microbiome-targeted and microbiome-friendly cosmetics has gained traction, with scientists investigating how specific microbial strains and metabolites can positively influence skin conditions such as acne, dryness, and inflammation (3). For example, *S. epidermidis*, a common skin commensal, has been shown to produce antimicrobial peptides that inhibit the growth of *S. aureus*, a pathogen linked to acne and eczema (4). This highlights the promise of incorporating beneficial strains into cosmetic formulations to help maintain or restore a balanced skin microbiome.

Microbiome cosmetics aim to either support the skin’s existing microbial community or introduce specific microbes or their byproducts to achieve targeted effects. Recent studies suggest that this approach could improve skin conditions in a more sustainable and biologically aligned way than traditional synthetic ingredients. Notably, ingredients derived from microbial sources, such as postbiotics (nonviable bacterial products or metabolic byproducts) and prebiotics (compounds that support beneficial microbes), are increasingly incorporated into skincare formulations for their ability to modulate skin pH, moisture, and immune responses (1).

The development of microbiome cosmetics is supported by growing evidence on the significance of a balanced skin microbiome for overall skin health and resilience against environmental stressors. Products that respect and promote microbial diversity on the skin may offer more personalized solutions by addressing specific skin needs through targeted microbial composition. This paradigm shift towards microbiome-based skincare highlights a promising growth area for the cosmetics industry, as consumers increasingly seek products that are both effective and biocompatible (5). *Corynebacterium* species have drawn attention to their influence on skin physiology, including hydration, immune modulation, and pigmentation. Recent research increasingly suggests that *Corynebacterium* species contribute not only to skin health maintenance but also to aesthetic properties, potentially through modulating factors. *Corynebacterium* species possess enzymes, such as lipases, that enable them to metabolize sebum, the skin’s natural oil, thereby helping maintain a balanced lipid layer on the skin surface (6–8). Furthermore, certain *Corynebacterium* species may have the potential to influence skin appearance, specifically by inhibiting melanin production. For example, *C. tuberculostearicum* produces a cyclic dipeptide, Cyclo (L-Pro-L-Tyr), which inhibits tyrosinase, a key enzyme in melanin synthesis. Tyrosinase inhibition has been linked to reduced melanin production, suggesting that *Corynebacterium* may influence skin brightness and tone, offering promising applications in cosmetic formulations aimed at skin brightening (8).

This study aims to explore the emerging field of microbiome cosmetics, focusing on bacterial postbiotics, their mechanisms of action on the skin, and their implications for future cosmetic product development.

## Results

### Bacterial isolation history and culture

In a previous study, skin microbiota samples were collected from the facial skin of a healthy female participant using sterile distilled water. A total of 100 colonies were obtained from TSB medium containing 0.1% Tween 80. 16S rRNA sequence analysis identified eight *Corynebacterium* strains (Table 1). All eight strains were successfully cultured in SMB medium.

**Table 1.**
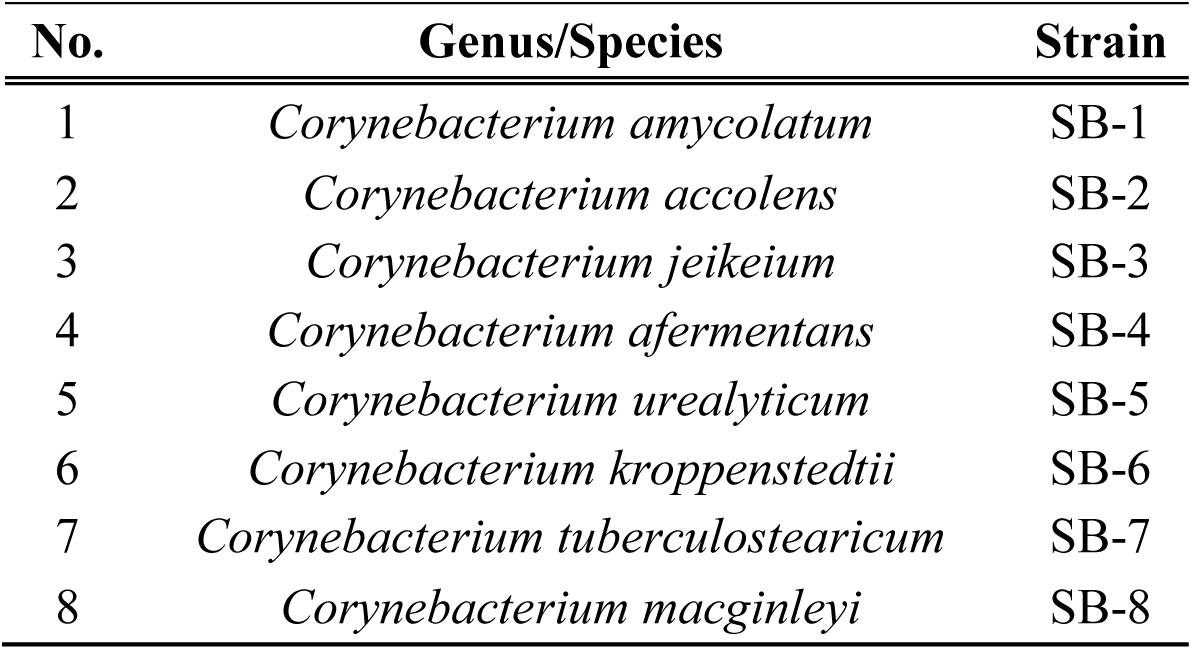
The list of isolated *Corynebacterium* genus.

### Skin-related effects of *Corynebacterium* ferment filtrates in human skin cell models

To compare the skin-related effects of the *Corynebacterium* ferment filtrates, we examined markers associated with extracellular matrix maintenance, inflammation, hydration, and pigmentation. The responses to the eight *Corynebacterium* isolates are summarized in Table 2. First, UVB-associated changes in extracellular matrix-related markers were evaluated in HS68 fibroblasts. Following ultraviolet B (UVB) exposure, significant decreases in elastin and *COL1A1* gene expression were observed. In contrast, *Corynebacterium* ferment filtrates (1–8) increased *ELN* expression, with strains 1 and 7 showing the largest increases and reaching levels above those of the non-UVB control (Fig. 1A). *Corynebacterium* ferment filtrates (1–7) also significantly increased *COL1A1* expression, with the largest responses observed for strains 1, 2, and 5 (Fig. 1B). In the case of the SB-8 strain, no positive effect was observed on the expression of the *COL1A1* gene.

**Figure 1.**
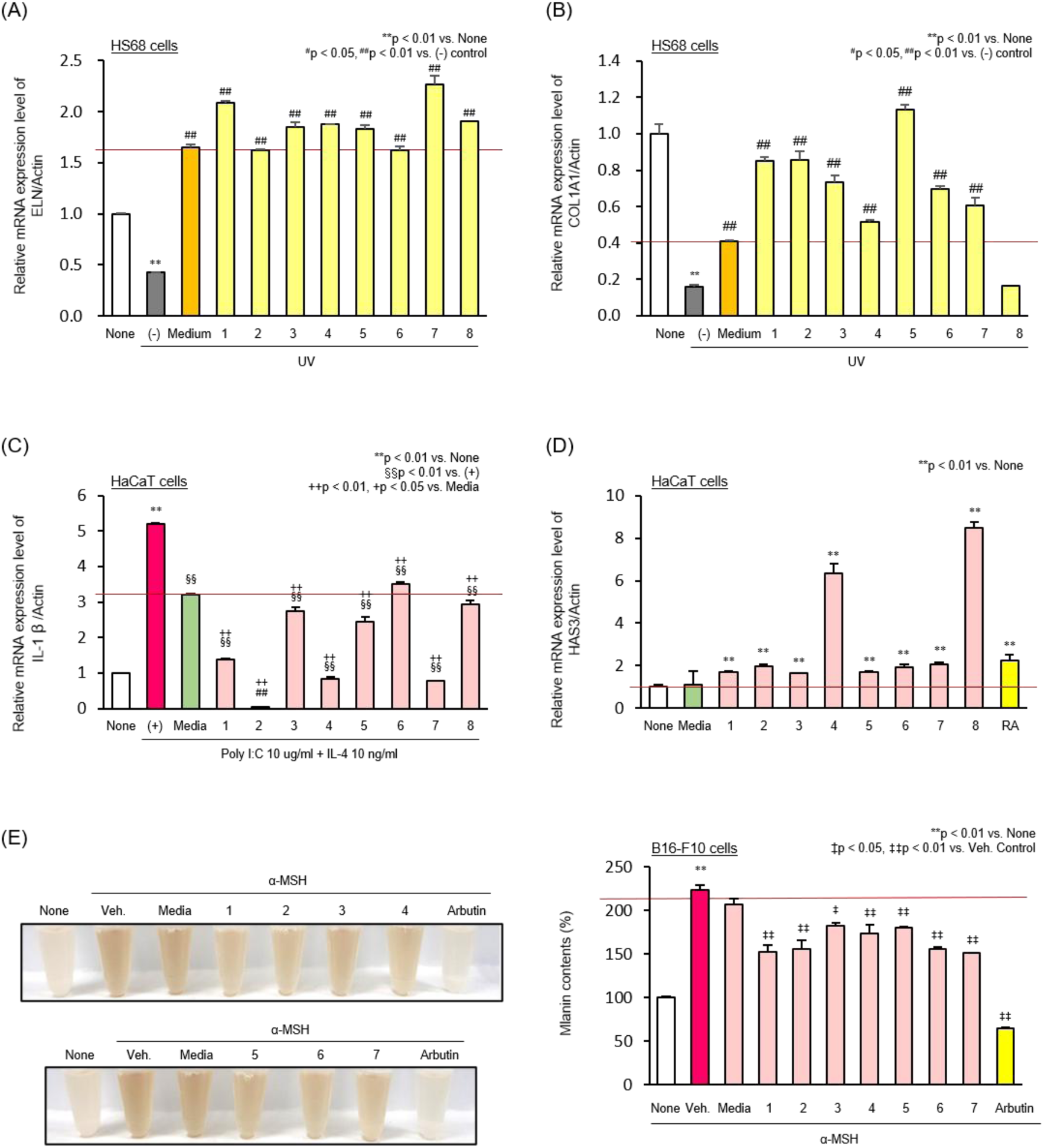
Effects of ferment filtrates from eight *Corynebacterium* strains in skin cell models. (A) Relative *ELN* expression in HS68 cells. (B) Relative *COL1A1* expression in HS68 cells. (C) Relative *IL-1β* expression in HaCaT cells. (D) Relative *HAS3* expression in HaCaT cells. (E) Relative melanin production. **p < 0.01 vs. None, ^#^p < 0.05, ^##^p < 0.01 vs. (−) control, §§ p < 0.01 vs. (+) control, ++p < 0.01, +p < 0.05 vs. Media, ‡p < 0.05, ‡‡p < 0.01 vs. Veh. Control.

**Table 2.** Skin condition improvement efficacy of 8 *Corynebacterium* strains.

| Strain | Elasticity |  | Anti-inflammation | Moisturization | Brightening | Product |
| --- | --- | --- | --- | --- | --- | --- |
| | Collagen | Elastin | IL-1β | HAS3 | Melanin | Growth |
| SB-1 | ++ | ++ | ++ | + | ++ | +++ |
| SB-2 | ++ | + | +++ | + | ++ | + |
| SB-3 | ++ | + | + | + | ++ | + |
| SB-4 | + | + | +++ | +++ | ++ | + |
| SB-5 | +++ | + | + | + | ++ | ++ |
| SB-6 | ++ | + | + | + | ++ | + |
| SB-7 | ++ | ++ | +++ | + | ++ | + |
| SB-8 |  | + | + | +++ |  | ++ |

The anti-inflammatory effects on *IL-1β* expression were evaluated in inflammation-induced HaCaT cells (Fig. 1C). Treatment with Poly I:C and *IL-1β* successfully induced inflammation, as evidenced by a significant increase in *IL-1β* expression compared to the normal group. Treatment with SMB medium alone resulted in a slight reduction in *IL-1β* expression but did not substantially mitigate the inflammatory response. In contrast, microbial culture filtrates (1–8) exhibited varying degrees of anti-inflammatory activity, with filtrates 1, 2, 4, and 7 showing the greatest reductions. These filtrates significantly reduced *IL-1β* expression to levels near or below that of the normal group.

The effects of the ferment filtrates on *HAS3* expression were evaluated in HaCaT cells. The normal group exhibited baseline *HAS3* expression, while treatment with SMB medium alone showed a minimal increase, indicating a limited effect of the medium itself. In contrast, microbial filtrates (1–8) demonstrated varying degrees of *HAS3* up-regulation, with filtrates 4 and 8 showing the most pronounced effects. These filtrates significantly increased *HAS3* expression, reaching levels comparable to or exceeding that of the positive control group treated with retinoic acid (Fig. 1D).

The inhibitory effect of microbial ferment filtrates on melanin production was assessed in α-melanocyte-stimulating hormone (α-MSH)-induced melanogenesis models. Normal cells exhibited baseline melanin levels, while α-MSH treatment significantly increased melanin production. Treatment with SMB medium alone showed no notable reduction in melanin levels, indicating that the medium itself had minimal effects. However, microbial culture filtrates (1–7) demonstrated varying degrees of melanin inhibition. Among these, filtrates 1, 2, 6, and 7 significantly reduced melanin production compared with the α-MSH-treated group, with values approaching those of the arbutin positive control (Fig. 1E). Based on the overall screening profile together with its growth characteristics, SB-1 was selected for further analysis.

### Safety Test of Corynebacterium amycolatum Strain SB-1

A hemolysis assay was conducted as an initial safety assessment of SB-1. The positive control, *S. aureus* ATCC 6538, showed clear β-hemolysis, whereas SB-1 exhibited no hemolytic activity (Fig 2A). Furthermore, the strain SB-1 was assessed for cytotoxicity by co-culturing it with HS68 and HaCaT cells. After 24 hours of incubation, cell viability was measured using the MTT assay. The results indicated that SB-1 did not reduce cell viability (Fig. 2B).

**Figure 2.**
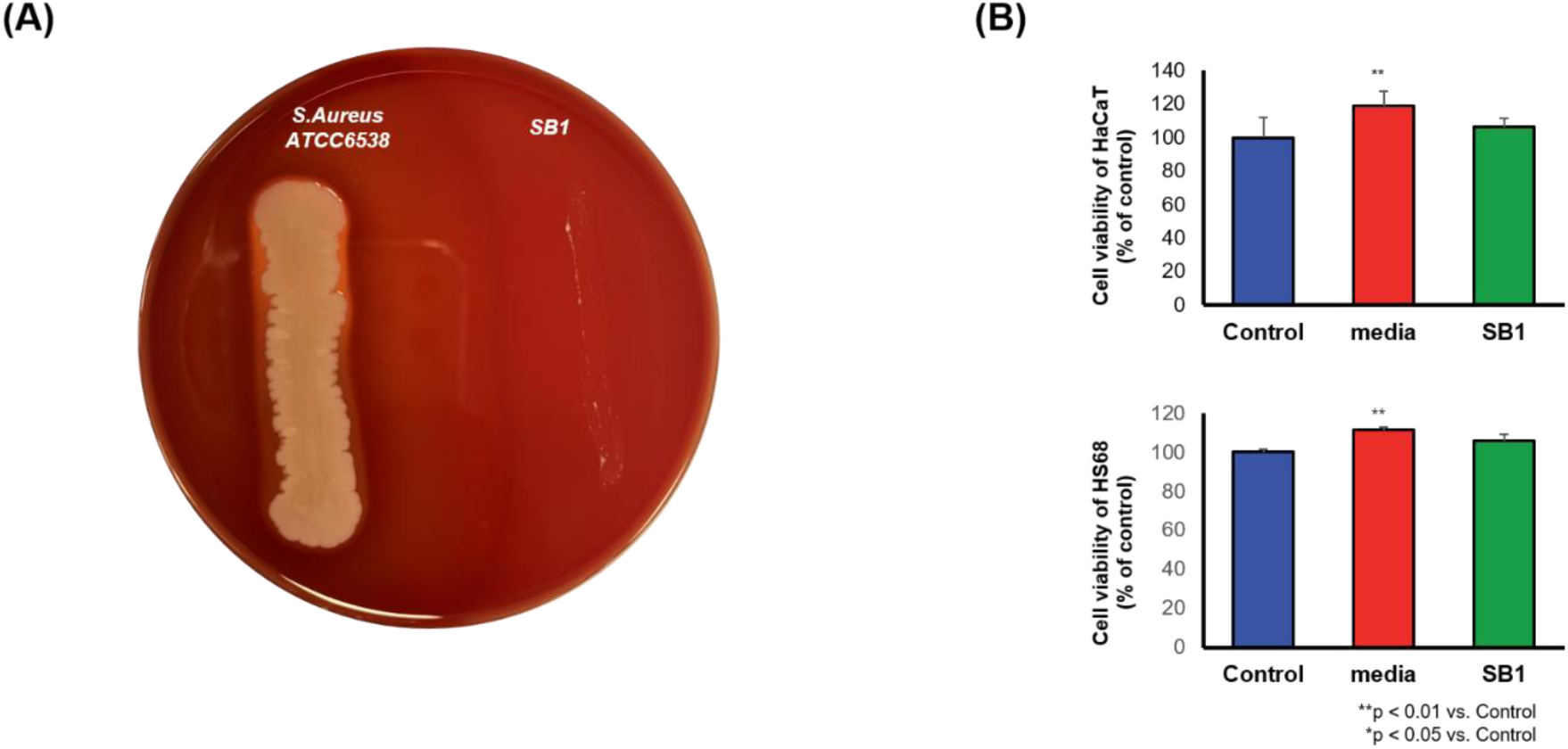
Safety test of *Corynebacterium amycolatum* strain SB-1. (A) Hemolytic test of strain SB-1. (B) Evaluation of cytotoxicity induced by SB-1 in HS68 and HaCaT Cells. **p < 0.01 vs. Control, *p < 0.05 vs. Control

### Screening and efficacy of ellagic acid identified in the SB-1 ferment filtrate

To screen for candidate metabolites, the ferment filtrate of strain SB-1 was analyzed using LC-TOF/MS. LC-TOF/MS analysis yielded an ellagic acid annotation at 48 ppm in the analytical dataset (Table 2; Fig. 3A). Pathway annotation was used to examine metabolic routes potentially related to ellagic acid or its precursors. A putative pathway considered in the present analysis involves both enzymatic and non-enzymatic processes, including routes from hydrolyzable tannins such as gallotannins and ellagitannins, as well as precursor formation via the shikimate and pentose phosphate pathways. Hydrolysable tannins are hydrolyzed to produce pentagalloylglucose, which further converts into hexahydroxydiphenic acid (HHDP) through hydrolysis and spontaneous lactonization. HHDP then undergoes spontaneous dimerization, esterification, or lactonization to form ellagic acid. Alternatively, gallic acid, a precursor for ellagic acid, can be synthesized from glucose-derived phosphoenolpyruvate and erythrose-4-phosphate through the shikimate pathway. Gallic acid combines glucose to form β-glucogallin, which subsequently transforms into ellagic acid. These annotations provide a putative metabolic context for the detected candidate; however, pathway annotation alone does not establish de novo ellagic acid biosynthesis by SB-1 (Fig. 3B).

**Figure 3.**
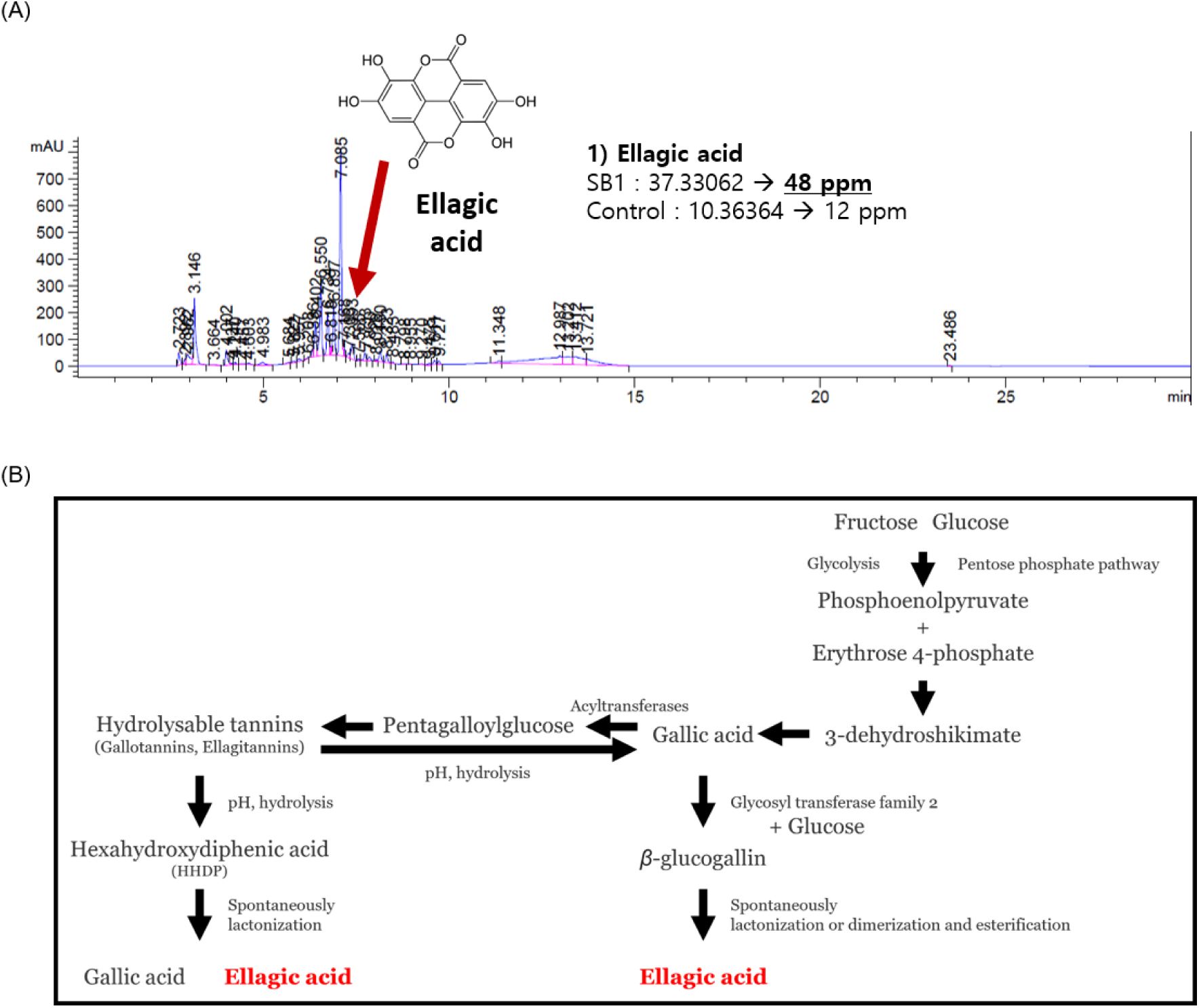
Screening of ellagic acid and a putative metabolic pathway in *Corynebacterium amycolatum* SB-1. (A) Candidate metabolite screening of the SB-1 ferment filtrate by LC-TOF/MS. (B) Putative metabolic pathway related to ellagic acid in *Corynebacterium amycolatum* SB-1.

To evaluate the effect of ellagic acid on a barrier-related marker, *FLG* expression was measured in HaCaT cells after treatment with ellagic acid (Table 2; Fig. 3B). In the normal control group, baseline levels of *FLG* expression were observed, and the dimethyl sulfoxide (100 nM of α-MSH + DMSO)-treated group showed no significant difference compared to normal cells. In contrast, treatment with retinoic acid (1μM), used as a positive control, resulted in a significant increase in *FLG* expression. Notably, *FLG* expression was significantly increased in cells treated with 4 μM ellagic acid, showing an increase in *FLG* transcript abundance under these conditions (Table 2; Fig. 4A).

**Figure 4.**
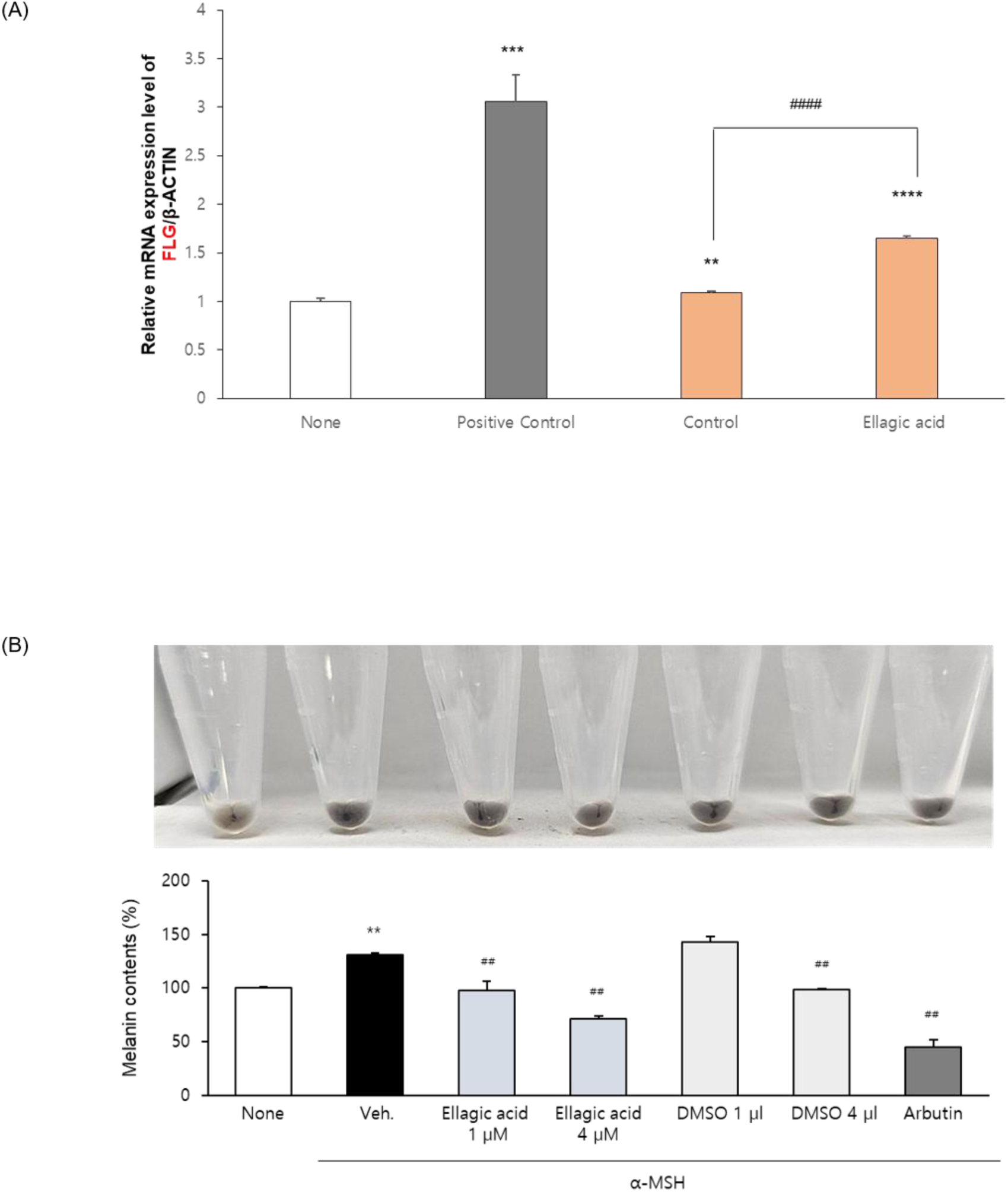
Effects of ellagic acid on *FLG* expression and melanin production. (A) Relative *FLG* expression following ellagic acid treatment. (B) Effect of ellagic acid on melanin production. All values are presented as the mean ± SD. **p < 0.01 vs. None, ##p < 0.01 vs. Veh.

In addition, the inhibitory effect of ellagic acid on melanin production was evaluated in α-MSH-induced melanogenesis model. Normal B16-F10 cells exhibited baseline melanin production, while α-MSH treatment significantly increased melanin production. Ellagic acid, dissolved in 100 nM of α-MSH + DMSO, reduced melanin production more than 100 nM of α-MSH + DMSO alone. These data show that ellagic acid reduced melanin content in this in vitro model (Table 2; Fig. 4B).

### Transcriptomic changes in RHPE following ellagic acid treatment

To investigate transcriptional changes, we analyzed RNA sequencing data from the Reconstructed Human Pigmented Epidermis model (RHPE) under four distinct conditions: the non-treated group, the 100 nM α-MSH-treated group, the 100 nM α-MSH with 100 nM of α-MSH + DMSO-treated group, and the 100 nM α-MSH with 4 µM ellagic acid-treated group. A total of 477,243,752 reads were generated, and the amount of uniquely mapped sequence per sample ranged from 4.36 to 6.00 Gb (Table 3). Principal Component Analysis revealed distinct clustering of samples by treatment conditions, indicating transcriptional differences among the experimental groups. The untreated group (3D-) formed a separate cluster, representing baseline transcriptional profiles, while the melanogenesis-induced group treated with α-MSH showed a distinct cluster associated with transcriptional changes induced by melanogenesis. The group treated with α-MSH and 100 nM of α-MSH + DMSO formed its own cluster, indicating that the vehicle contributed to transcriptional variation, whereas the group treated with α-MSH and ellagic acid also clustered separately. Thus, the PCA demonstrates treatment-associated separation but does not by itself distinguish ellagic acid-specific effects from the 100 nM of α-MSH + DMSO vehicle effect (Figure 5A).

**Figure 5.**
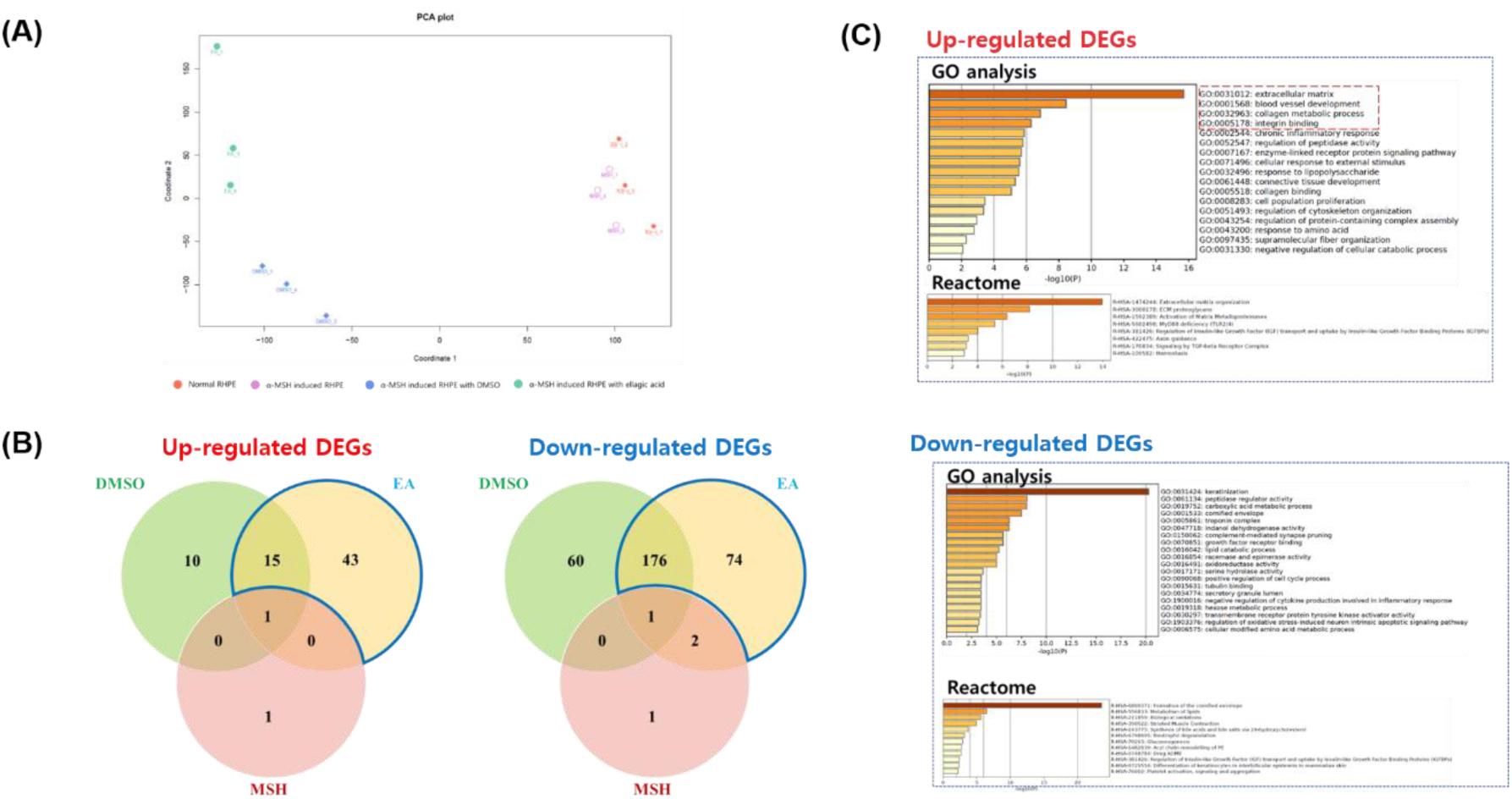
Transcriptomic profiling and functional enrichment analysis of ellagic acid treatment. (A) Principal component analysis of RNA-seq data showing sample-to-sample variation across the four groups (the non-treated group, 100 nM of α-MSH treated group, 100 nM of α-MSH and 100 nM of α-MSH + DMSO treated group, and 100 nM of α-MSH and 4 µM ellagic acid-treated group). (B) Venn diagrams illustrating the unique and overlapping up-regulated and down-regulated DEGs in the α-MSH (100 nM), α-MSH with 100 nM of α-MSH + DMSO, and α-MSH with ellagic acid (4 µM) groups compared to the untreated control. (C) Functional enrichment analysis of DEG sets associated with the ellagic-acid-containing group.

**Table 3.** Summary of sequencing statistics.

| **Treatment** | **Total reads** | **Unique mapped reads Count** | **%** | **Multiple mapped reads Count** | **%** | **Unmapped reads Count** | **%** |
| --- | --- | --- | --- | --- | --- | --- | --- |
| 3D(-)_1 | 36,925,832 | 32,351,863 | 87.61% | 3,212,946 | 8.70% | 1,279,279 | 3.46% |
| 3D(-)_2 | 37,694,036 | 33,835,278 | 89.76% | 3,189,596 | 8.46% | 572,737 | 1.52% |
| 3D(-)_3 | 38,122,442 | 33,844,098 | 88.78% | 3,459,374 | 9.07% | 728,228 | 1.91% |
| MSH_1 | 44,545,705 | 40,027,863 | 89.86% | 2,892,611 | 6.49% | 1,538,681 | 3.45% |
| MSH_2 | 44,598,480 | 39,041,573 | 87.54% | 3,463,448 | 7.77% | 1,994,127 | 4.47% |
| MSH_4 | 43,052,941 | 37,918,891 | 88.08% | 3,379,053 | 7.85% | 1,661,037 | 3.86% |
| DMSO_1 | 37,985,061 | 34,499,127 | 90.82% | 2,787,313 | 7.34% | 607,648 | 1.60% |
| DMSO_3 | 44,652,693 | 39,699,787 | 88.91% | 3,825,628 | 8.57% | 1,028,378 | 2.30% |
| DMSO_4 | 33,295,631 | 29,127,552 | 87.48% | 2,784,103 | 8.36% | 1,305,310 | 3.92% |
| EA_1 | 32,166,573 | 29,099,792 | 90.47% | 2,324,201 | 7.23% | 679,990 | 2.11% |
| EA_3 | 39,799,557 | 35,100,082 | 88.19% | 3,185,541 | 8.00% | 1,425,959 | 3.58% |
| EA_4 | 44,404,801 | 39,714,598 | 89.44% | 3,450,314 | 7.77% | 1,153,201 | 2.60% |

Differentially expressed gene (DEG) analysis was performed using the stated cutoff criteria (adjusted p-value < 0.05 and log2 FC ≥ 1) to identify significant transcriptional changes across the experimental conditions. The table summarizes the number of DEGs for each comparison, highlighting the transcriptional impact of the treatments (Table 4).

**Table 4.** Differentially Expressed Gene (DEG) analysis.

| Comparison | Condition | Up-regulation | Down-regulation | Total |
| --- | --- | --- | --- | --- |
| 3D(-) vs α-MSH | 2-fold | 153 | 433 | 586 |
|  | 2-fold & p<0.05 | 2 | 4 | 6 |
| 3D(-) vs DMSO | 2-fold | 436 | 1131 | 1567 |
|  | 2-fold & p<0.05 | 26 | 237 | 263 |
| 3D(-) vs Ellagic acid | 2-fold | 418 | 1535 | 1953 |
|  | 2-fold & p<0.05 | 59 | 253 | 312 |

As shown in Figure 5, the DEG results showed changes in a total of six genes in the α-MSH-treated group. Of these, two genes showed up-regulated and four genes showed down-regulated expression compared with the normal group. The 100 nM of α-MSH + DMSO-treated group displayed a total of 263 differentially expressed genes. Of these, 26 genes showed up-regulated expression, and 237 showed down-regulated expression compared to the normal group, reflecting significant alterations in gene expression. The ellagic acid-treated group demonstrated a total of 312 differentially expressed genes. Among these, 59 genes were found to be up-regulated, while 253 genes were down-regulated compared to the normal control group, indicating significant changes in the expression of these genes (Fig. 5). These results show that the 100 nM of α-MSH + DMSO-containing groups had substantially more DEGs relative to untreated control than the α-MSH-only group. Because 100 nM of Α-MSH + DMSO alone was associated with 263 DEGs, a direct ellagic acid-versus-vehicle comparison is required to define the ellagic acid-specific component of the response.

Based on the Venn diagram, a total of 43 genes were identified as uniquely up-regulated in the ellagic-acid-containing group relative to the untreated-control-based DEG sets. Functional enrichment analyses revealed that these up-regulated genes are primarily involved in biological processes such as extracellular matrix organization, blood vessel development, and collagen metabolic processes. Additionally, these genes were enriched in molecular functions related to integrin binding, indicating enrichment of extracellular matrix- and cell adhesion-related transcriptional programs in this gene set (Fig. 5). A total of 43 genes were also identified as uniquely down-regulated in the ellagic-acid-containing group relative to the untreated-control-based DEG sets. GO analysis revealed that these down-regulated genes are associated with biological processes such as keratinization, peptidase regulator activity, and carboxylic acid metabolic processes. Furthermore, Reactome pathway analysis indicated that these genes are involved in pathways related to the formation of the cornified envelope, lipid metabolism, and biological oxidation. These results describe pathways represented among genes unique to the ellagic-acid-containing group; because the DEG sets were defined relative to the untreated control, they should not be interpreted as vehicle-adjusted ellagic acid effects (Fig. 5).

Heatmap analysis of pigment granule organization-related genes showed a distinct expression pattern in the ellagic-acid-containing group for genes associated with melanin synthesis and granule transport. Notably, the *TYRP1* gene, critical for melanin biosynthesis, was significantly down-regulated, consistent with lower TYRP1 transcript abundance. Furthermore, *RAB32* and *RAB27A*, which play key roles in melanosome transport and docking, were also down-regulated, indicating lower expression of genes involved in melanosome transport and docking. Similarly, *BLOC1S1* and *HPS4*, genes involved in the biogenesis of lysosome-related organelles, showed decreased expression. In contrast, several melanosome organization-related genes showed higher expression in the ellagic-acid-containing group. Among the selected genes, *MITF*, a master regulator of melanogenesis, was significantly up-regulated, showing that the response was not uniformly suppressive across the melanogenesis program. Similarly, *PMEL*, which is crucial for the structural organization of melanosomes, and *BCL2A1*, involved in cell survival and melanosome formation, were also up-regulated in the ellagic-acid-containing group. Furthermore, *GPR143*, a gene essential for melanosome transport and signaling, showed increased expression, indicating higher *GPR143* transcript abundance. On the other hand, *ADAMTSL4*, which is involved in extracellular matrix interactions, was uniquely down-regulated, showing lower *ADAMTSL4* expression. Together, these mixed directions indicate remodeling of pigmentation- and melanosome-related transcription rather than uniform suppression of the pathway (Fig. 6A).

**Figure 6.**
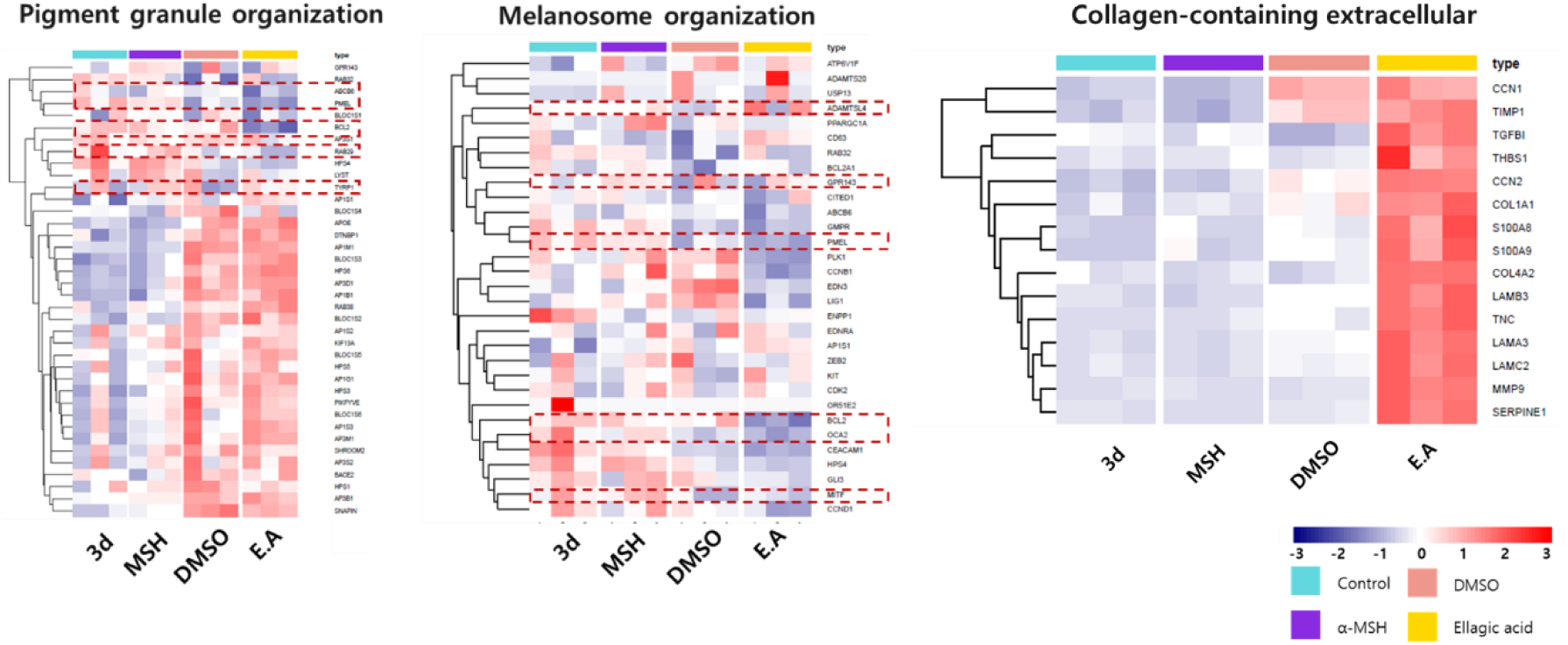
Expression profiles of key genes involved in pigmentation and extracellular matrix organization. Hierarchical clustering heatmaps illustrating the expression patterns of DEGs related to (A) pigment granule organization, (B) melanosome organization, and (C) the collagen-containing extracellular matrix across the four experimental groups (non-treated, 100 nM α-MSH, 100 nM α-MSH with 100 nM of α-MSH + DMSO, and 100 nM α-MSH with 4 µM ellagic acid). The color scale represents the normalized expression levels.

Genes associated with the collagen-containing extracellular matrix showed transcriptional changes in the ellagic-acid-containing group. Among the selected genes, *CCN2*, a key regulator of extracellular matrix remodeling and fibrosis, was notably up-regulated, indicating higher *CCN2* transcript abundance. Similarly, *COL1A1* and *COL4A2*, critical genes encoding type I and IV collagen, respectively, were significantly up-regulated, indicating higher transcript levels of type I- and IV-collagen genes. *LAMC2*, a gene involved in laminin complex assembly, was also up-regulated, indicating higher *LAMC2* transcript abundance. Additionally, *MMP9*, a matrix metalloproteinase involved in collagen degradation, and *SERPINE1*, a serine protease inhibitor that regulates extracellular matrix turnover, were uniquely up-regulated, indicating coordinated changes in genes involved in extracellular matrix turnover and remodeling (Fig. 6B).

## Discussion

The successful isolation of eight *Corynebacterium* isolates in this study was achieved using TSB medium supplemented with 0.1% Tween 80. Tween 80, a nonionic surfactant, is known to enhance the growth of lipophilic *Corynebacterium* species by providing essential fatty acids and reducing surface tension, thereby facilitating better nutrient absorption. This approach aligns with previous findings that demonstrate the necessity of lipid supplementation for the optimal cultivation of certain *Corynebacterium* species (9, 10).

The evaluation of cosmetic ingredients’ efficacy on skin improvement is often conducted through in vitro cell-based assays that assess the expression of genes representative of skin characteristics. This approach allows for the identification of active compounds that can modulate specific skin functions, providing a useful screening approach before more direct functional or in vivo validation (11).

In our study, we focused on several key genes associated with skin elasticity, inflammation, hydration, barrier function, and pigmentation (Fig.1A and 1B). The expression levels of *COL1A1* and *ELN* were analyzed because their encoded proteins are major extracellular matrix components that contribute to skin firmness and elasticity. Previous research has demonstrated that collagen peptides can enhance the expression of these genes, leading to improved skin elasticity and firmness (12, 13). Ultraviolet (UV) radiation is a well-established factor contributing to skin aging, primarily through the degradation of extracellular matrix components such as elastin and collagen type I (*COL1A1*). This degradation leads to reduced skin elasticity and the formation of wrinkles. In our study, UV treatment significantly down-regulated the expression of elastin and *COL1A1* in HS68 fibroblast cells, corroborating previous findings on UV-induced dermal damage.

Notably, treatment with *Corynebacterium* ferment filtrates (strains 1–8) increased the expression of these extracellular matrix-related transcripts. Strains 1 and 7 not only restored but also enhanced elastin expression beyond normal levels, while strains 1, 2, and 5 significantly increased *COL1A1* expression (Fig.1A and Fig.1B). These results suggest that selected *Corynebacterium* ferment filtrates can partially restore UVB-associated reductions in extracellular matrix-related transcripts, supporting further evaluation of their effects on dermal matrix homeostasis. The beneficial effects of microbial ferment filtrates on skin health have been documented in previous studies. For instance, a study on *Lactobacillus plantarum* GT-17F fermented Dendrobium officinale extract demonstrated significant improvements in skin elasticity and wrinkle reduction in human subjects after 28 days of treatment (14). Similarly, topical application of *Epidermidibacterium keratini* EPI-7 ferment filtrate resulted in enhanced skin elasticity and dermal density, as evidenced by a randomized split-face clinical study (14). These findings align with our observations, indicating that microbial ferment filtrates can modulate gene expression related to skin structure and function. The observed changes in *ELN* and *COL1A1* provide a basis for further functional testing, although collagen and elastin protein abundance were not measured in the present study. Further research, including clinical trials, is warranted to assess the efficacy and safety of these fermented filtrates for cosmetic applications.

Additionally, we examined the expression of interleukin-1 beta (*IL-1β*), a pro-inflammatory cytokine that plays a pivotal role in skin inflammation (Fig. 1C). Elevated levels of *IL-1β* are associated with various inflammatory skin conditions, making it a valuable marker for assessing the anti-inflammatory potential of cosmetic ingredients (15). Treatment with SMB medium alone led to a slight reduction in *IL-1β* levels but did not substantially mitigate the inflammatory response. In contrast, microbial culture filtrates (1–8) exhibited varying degrees of anti-inflammatory activity, with filtrates 1, 2, 4, and 7 demonstrating the highest efficacy, significantly reducing *IL-1β* expression to levels near or below those of the normal group. These findings show that selected microbial ferment filtrates reduced *IL-1β* expression in this in vitro model, supporting further evaluation of their anti-inflammatory potential. This aligns with previous research indicating that certain microbial-derived products can modulate inflammatory pathways in skin cells. For instance, a study on the effects of a lotion containing probiotic ferment lysate demonstrated enhanced skin barrier function and reduced inflammation in vitro and in vivo models (14). Because chronic inflammation can affect extracellular matrix homeostasis, attenuation of inflammatory signaling may be relevant to structural skin maintenance; however, this relationship was not directly tested in the present study.

To assess skin hydration and barrier function, we evaluated the expression of filaggrin (*FLG*) and hyaluronan synthase 3 (*HAS3*) (Fig.1D). Filaggrin is essential for the formation of the skin barrier and maintaining moisture levels, while *HAS3* is involved in the synthesis of hyaluronic acid (HA), a key molecule in skin hydration. Alterations in the expression of these genes can significantly impact skin moisture retention and barrier integrity (15, 16). Selected microbial ferment filtrates increased *HAS3* transcript expression; whether this translated into greater hyaluronic acid production was not directly measured. This aligns with previous research demonstrating the beneficial effects of microbial-derived products on skin health. For instance, a study on the effects of a lotion containing probiotic ferment lysate reported enhanced skin barrier function and increased skin hydration, attributed to the up-regulation of genes involved in skin barrier and hydration processes (17). *HAS3* up-regulation is biologically relevant because increased hyaluronan synthesis can promote water retention, and previous studies have linked microbial-derived products to hydration-related responses. Nevertheless, HA content, hydration, and elasticity were not directly measured in this study. Taken together, the data support a transcriptional effect of selected ferment filtrates on *HAS3* in HaCaT cells and justify direct measurement of HA production and barrier or hydration endpoints in future studies.

Furthermore, we investigated genes related to melanin production, as melanin is the primary determinant of skin tone and brightness (Fig.1E). Modulation of melanin synthesis pathways can lead to changes in pigmentation, which are crucial for cosmetic applications aimed at skin lightening or evening out skin tone (18, 19). By analyzing the expression patterns of these genes in response to cosmetic ingredients, we can infer their potential effects on skin properties such as elasticity, inflammation, hydration, barrier function, and pigmentation. This cell-based approach provides a valuable tool for screening and developing cosmetic products with targeted skin benefits. Notably, treatment with microbial culture filtrates (1–7) resulted in varying degrees of melanin inhibition, with filtrates 1, 2, 6, and 7 demonstrating significant reductions in melanin levels, comparable to the effects observed with arbutin, a well-known melanin inhibitor. These findings show that selected microbial ferment filtrates reduced melanin production in this in vitro screening model and may be relevant to cosmetic pigmentation control (8). These results support further evaluation of selected ferment filtrates as pigmentation-modulating cosmetic candidates. No direct relationship between melanin reduction and skin elasticity was assessed in this study.

Evaluating the safety of microbial strains considered for cosmetic applications is important, particularly concerning their hemolytic activity and cytotoxicity. In our study, strain SB-1 exhibited no hemolytic activity, as evidenced by the absence of hemolysis in hemolysis testing, whereas the positive control, *S. aureus* ATCC 6538, displayed clear beta-hemolysis (Fig. 2A) (20). Furthermore, the MTT assay results demonstrated that co-culturing strain SB-1 with HS68 and HaCaT cells for 24 hours did not reduce cell viability, indicating a lack of cytotoxic effects (Fig. 2B). This outcome is consistent with studies assessing the cytotoxicity of probiotic strains, where non-pathogenic strains did not adversely affect cell viability (21). Together, these assays provide initial evidence that SB-1 is non-hemolytic and did not reduce cell viability under the tested co-culture conditions. These assays do not constitute a comprehensive safety assessment, and additional studies are required before use in cosmetic formulations.

Ellagic acid, a naturally occurring polyphenol, has garnered attention for its potential skin benefits, particularly in enhancing skin barrier function and reducing hyperpigmentation (22–24). In our study, ellagic acid treatment significantly up-regulated *FLG* mRNA expression in keratinocytes, surpassing the effects observed with the positive control, retinoic acid (Fig. 4A). Filaggrin is a crucial protein in the epidermis, essential for skin barrier integrity and hydration. Its up-regulation is consistent with a possible effect on barrier-related differentiation; however, barrier function itself was not directly measured. Additionally, ellagic acid demonstrated a pronounced inhibitory effect on melanin production in an α-MSH-induced melanogenesis model. This reduction in melanin production supports further evaluation of ellagic acid as a pigmentation-modulating candidate in cosmetic models. These findings are consistent with previous research highlighting ellagic acid’s role in skin health.

RNA sequencing of the RHPE model showed treatment-associated transcriptional differences under melanogenesis-induced conditions. Principal Component Analysis revealed that the transcriptional profile of the ellagic acid -treated group was distinctly separated from all other groups, indicating a distinct expression profile (Figure 5A). Because the ellagic acid condition also contained 100 nM of α-MSH + DMSO, PCA separation alone cannot establish which component produced the observed differences (22). It is noteworthy that α-MSH was utilized to induce melanogenesis in this study. However, the transcriptional differences between the normal RHPE cells 3D (−) and the α-MSH-treated group were relatively subtle. One possible explanation is the intrinsic phenotype of the phototype VI RHPE model, which has a higher basal level of pigmentation than lighter phototypes. Consequently, the normal RHPE cells already displayed transcriptional features associated with melanogenesis, reducing the observable transcriptional shifts upon α-MSH stimulation (22).

Differential gene expression analysis showed marked differences across the experimental groups, with the two 100 nM of α-MSH + DMSO-containing groups showing substantially more DEGs relative to untreated control than α-MSH alone. The α-MSH-treated group exhibited minimal transcriptional alterations, with only 6 DEGs identified. Among these, 2 genes were up-regulated, and 4 genes were down-regulated, reflecting the limited transcriptional shift induced by α-MSH. In contrast, the 100 nM of α-MSH + DMSO-treated group displayed significant transcriptional changes, with 263 DEGs identified. Of these, 26 genes were up-regulated, and 237 genes were down-regulated. This suggests that 100 nM of α-MSH + DMSO, despite its common use as a solvent, exerts notable transcriptional regulatory effects. Previous studies have reported 100 nM of α-MSH + DMSO’s ability to modulate cellular pathways, including those related to oxidative stress and inflammation, which may partially explain these observations (25). The ellagic acid-treated group exhibited the largest transcriptional impact, with 312 DEGs, including 59 up-regulated and 253 down-regulated genes. However, because the 100 nM of α-MSH + DMSO vehicle alone was associated with 263 DEGs, the additional differences observed in the ellagic-acid-containing group cannot be attributed specifically to ellagic acid from untreated-control comparisons alone. A direct comparison between α-MSH plus ellagic acid and α-MSH plus 100 nM of α-MSH + DMSO is therefore required to define the vehicle-adjusted ellagic acid response. Interestingly, microbial ferment extracts have also demonstrated the ability to influence RNA-level changes in skin cells, supporting the idea that external treatments can modulate gene expression in the skin. Previous studies have shown that bioactive compounds from microbial fermentation can alter pathways associated with skin barrier function, oxidative stress, and pigmentation, further highlighting the dynamic responsiveness of skin cells to external bioactive agents (26, 27).

The ellagic-acid-containing group showed transcriptional differences in pigmentation- and extracellular matrix (ECM)-related genes, as evidenced by Venn diagram and heatmap analyses. Venn diagram analysis identified 43 uniquely upregulated genes, which were primarily associated with biological processes such as extracellular matrix organization, blood vessel development, and collagen metabolic processes. GO and Reactome pathway analyses further highlighted the enrichment of these genes in molecular functions like integrin binding, indicating enrichment of ECM- and cell adhesion-related transcriptional programs. Key genes such as *CCN2*, a central regulator of ECM remodeling, were significantly upregulated, alongside *COL1A1* and *COL4A2*, encoding type I and IV collagen, respectively, showing higher transcript abundance of structural matrix genes. Up-regulation of *LAMC2*, which supports basement membrane assembly, further indicates higher expression of a basement membrane-related gene. Concurrently, *MMP9* and *SERPINE1*, involved in ECM turnover and remodeling, were also upregulated, showing coordinated transcriptional changes in genes associated with ECM synthesis and turnover (22, 24).

Heatmap analysis showed changes in genes associated with melanin synthesis, pigment granule transport, and melanosome organization in the ellagic-acid-containing group. Down-regulation of *TYRP1*, a critical enzyme in melanin biosynthesis, alongside *RAB32* and *RAB27A*, which are essential for melanosome transport and docking, was consistent with lower expression of several genes involved in pigment synthesis and granule trafficking. Similarly, genes like *BLOC1S1* and *HPS4*, involved in the biogenesis of lysosome-related organelles, were down-regulated, further showing lower expression of genes associated with melanosome maturation. Interestingly, ellagic acid also up-regulated key regulators of melanosome biogenesis, such as *MITF*, the master transcription factor of melanogenesis, and *PMEL*, which contributes to melanosome structural integrity. The increased expression of *BCL2A1*, promoting cell survival and melanosome formation, and *GPR143*, involved in melanosome transport and signaling, demonstrates that the transcriptional response was bidirectional rather than a uniform inhibition of melanogenesis. Additionally, down-regulation of *ADAMTSL4*, associated with ECM interactions, adds to this mixed expression pattern and warrants further mechanistic evaluation.

Together, these findings support further study of ellagic acid in skin-related in vitro models. The transcriptomic data indicate changes in pigmentation- and ECM-related programs, but vehicle-controlled differential expression analysis and direct functional validation are needed before assigning ellagic acid-specific mechanisms or clinical relevance.

## Materials and Methods

### Isolation and identification of the interested *Corynebacterium* Strains

Skin microbiota samples were collected from the facial skin of a healthy female participant by washing the designated area with sterile distilled water. The samples were inoculated into a liquid medium containing tryptic soy broth (TSB) (Becton Dickinson, Franklin Lakes, New Jersey, USA) with 0.1% Tween 80, or onto a tryptic soy agar (TSA) solid medium (Becton Dickinson, Franklin Lakes, New Jersey, USA). Following incubation at 28°C for 48 hours, 100 colonies were isolated and subjected to pure culture, then incubated for an additional 48 hours at 28°C. The colonies were identified through 16S rRNA gene sequencing, using primers specifically designed for bacterial DNA amplification (27F: 5’-AGAGTTTGATCCTGGCTCAG-3’ and 1492R: 5’-GGTTACCTTGTTACGACTT-3’). PCR conditions included 30 cycles at 95°C for 1 minute, 55°C for 1 minute, and 75°C for 1 minute 30 seconds, followed by a final extension at 72°C for 8 minutes. Samples were then stored at 4°C. The DNA sequences of the isolated strains were determined using an ABI-3730XL sequencer (ABI, Waltham, Massachusetts, USA), and sequence analysis was conducted using the BLAST program on the EzBioCloud website (28). The genome sequencing data for *Corynebacterium amycolatum* strain (SB-1) are accessible from NCBI Database (Accession number: CP120206.1 and Bioproject: PRJNA943520).

### Preparation of *Corynebacterium* culture filtrates

Eight *Corynebacterium* isolates were cultured in TSB or on TSA at 30 °C for 48 h. A single colony was picked from the plate and grown overnight in a customized skin-mimetic broth (SMB). The SMB was made of 1.06 g of glucose, 0.2 g of rhamnose, 0.013 g of fructose, 0.05 g of MgSO4, 2.5 g of K2HPO4, 5 g of NaCl, and 1 g of yeast extract in a liter of distilled water. Fifty milliliters of fresh SMB were inoculated with the overnight culture at a 1:100 dilution, transferred to a 250-mL Erlenmeyer flask, and incubated at 30 °C with shaking at 160 rpm for 48 h in the dark. The culture was filtered with a syringe filter (0.45-μm pore size; Minisart, Sartorius, Göttingen, Germany). All culture filtrates were prepared in triplicate (16).

### Skin Cell Culture

To evaluate the effects of culture supernatants from *Corynebacterium* strains on the skin, we used three cell lines: human dermal fibroblasts (HS68), human keratinocytes (HaCaT), and murine melanoma cells (B16-F10). The Hs68, HaCaT, and B16-F10 cell lines, which is derived from C57BL/6J murine skin cells, were purchased from American Type Culture Collection (ATCC; Manassas, VA, USA). Cells were cultured in Dulbecco’s Modified Eagle’s Medium (DMEM; Gibco 1210-0038) supplemented with 10% fetal bovine serum (FBS) (Thermo Fisher Scientific, Waltham, Massachusetts, USA) at 37°C in a 5% CO₂ atmosphere. The culture medium was refreshed every 3–4 days, and cells were subcultured upon reaching confluence. For experiments, cells were seeded at a density of 5×10⁵ cells per well in appropriate culture plates. After a 24-hour incubation period, cells were washed with phosphate-buffered saline (PBS).

### Efficacy Test

To assess UVB-associated changes in extracellular matrix-related gene expression, HS68 cells were exposed to 12 mJ/cm² of UVB radiation, followed by a 24-hour incubation with 1% (w/w) SB-1 supernatant in a serum-free medium. Negative control groups included cells neither exposed to UVB nor treated with bacterial culture supernatants.

HaCaT cells were used to evaluate inflammation-related, skin barrier-related, and hydration-related responses. Inflammation was induced by treating the cells with 10 μg/mL polyinosinic– polycytidylic acid (poly I:C) and 10 ng/mL interleukin-1β (*IL-1β*). Subsequently, 1% (w/w) SB-1 supernatant was added, and the cells were incubated for 4 hours. In addition, the effects of ellagic acid (Sigma-Aldrich, Burlington, Massachusetts, USA) on skin barrier function and moisturizing activity were evaluated. The cells were treated with 4µM of ellagic acid and incubated for an additional 24 hours. Retinoic acid at a concentration of 1μM was used as the positive control.

To investigate melanin synthesis, the B16-F10 cell line (1×10⁵ cells/well) was stimulated with 0.1μM α-MSH and treated with 1% SB-1 for 72 hours. Arbutin (100 ppm; Sigma-Aldrich), known for its inhibitory effect on melanogenesis, was used as a positive control. Post-incubation, the cells were harvested and the cell pellet (1×10⁶ cells/mL) was collected. The cell pellets were dried at 60°C for 1 hour and then lysed in 1M NaOH containing 10% dimethyl sulfoxide (100 nM of α-MSH + DMSO) to solubilize the intracellular melanin. The quantity of melanin was measured by determining the absorbances at 490 nm using a microplate reader Victor3 (PerkinElmer, Shelton, Connecticut, USA) (18). The inhibition of melanin production was calculated as a percentage of the control using the following formula: Melanin content (%) = (Absorbance of control group/Absorbance of treated group) × 100.

### Hemolysis assay

A hemolysis assay was performed using blood agar to evaluate the hemolytic activity of the strain SB-1. Blood agar plates were prepared by supplementing a sterile agar base medium with 5% defibrinated sheep blood. SB-1 and the positive control S. aureus ATCC 6538 were streaked onto blood agar plates using a sterile inoculating loop. The plates were then incubated at 30°C for 48 hours under aerobic conditions. After incubation, the plates were examined for hemolytic activity, which was categorized as alpha (α) hemolysis, indicating partial hemolysis with a greenish discoloration; beta (β) hemolysis, representing complete hemolysis with a clear zone; or gamma (γ) hemolysis, indicating no hemolysis (29).

### MTT assay

To evaluate the cytotoxicity of the SB-1 strain, HaCaT and HS68 cells were co-cultured for 24 hours using CellQART co-culture wells (SABEU, Ennepetal, North Rhine-Westphalia, Germany) under standard conditions (37°C in a humidified atmosphere with 5% CO₂). After incubation, cell viability was assessed using the MTT assay. The culture medium was replaced with fresh medium containing MTT solution (final concentration of 0.5 mg/mL), and the plates were incubated again at 37°C with 5% CO₂ for 3–4 hours to allow viable cells to reduce MTT to formazan crystals. Following incubation, the medium was carefully removed, and 100 nM of α-MSH + DMSO was added to dissolve the formazan crystals. Absorbance was measured at 570 nm with a reference wavelength of 650 nm using a microplate reader. Cell viability was calculated as a percentage relative to untreated control cells, based on the absorbance values. This method followed the standardized MTT assay protocol.

### Quantitative PCR

RNA extraction from each sample was performed using TRI reagent (Takara, Tokyo, Kantō, Japan), and RNA concentration was measured using a NanoDrop spectrophotometer. For cDNA synthesis, 2 μg of RNA was reverse-transcribed using a thermal cycler (C1000, Bio-Rad, USA). Quantitative real-time PCR was conducted to amplify collagen type I alpha 1 chain (*COL1A1*), elastin (*ELN*), pro-inflammatory cytokine gene interleukin-1 beta (*IL-1β*), filaggrin (*FLG*), and hyaluronan synthase 3 (*HAS3*), employing SYBR Green Supermix (Applied Biosystems, Waltham, Massachusetts, USA) and specific primers on a StepOnePlus Real-Time PCR System (Applied Biosystems, Waltham, Massachusetts, USA). Gene expression levels were normalized to β-actin and analyzed accordingly (16).

### Metabolomic Analysis

For metabolomic analysis, **s**train SB-1 was inoculated into 1 L of SMB medium and cultured at 30°C with shaking at 160 rpm for 48 hours. The culture broth was then centrifuged at 6,000 rpm for 20 minutes to separate the cells. The supernatant was collected and filtered through a syringe filter with a 0.45-μm pore size (Minisart, Sartorius, Göttingen, Germany) to remove any remaining cells. The filtered supernatant was subsequently freeze-dried at −80°C, and the lyophilized sample was collected for further analysis.

The 1290 Infinity II UHPLC and 6545XT AdvanceBio Q-TOF Systems (Agilent Technologies, Santa Clara, California, USA), as well as the InfinityLab Poroshell 120 EC-C18 Column packed (2.7-µm, 2.1 × 150 mm, Agilent Technologies, Santa Clara, California, USA), were used for metabolomic analysis. The mobile phase comprised water with 0.1% formic acid (A) and acetonitrile with 0.1% formic acid (B), and chromatographic separation was performed at 40 °C. The chromatographic gradient was set as follows: 0–10 min, 0–40% B; 10–18 min, 40–95%; 18–27 min, 95% B; 27–27.5 min, 95–0% B; and 27.5–30 min, 0% B. Five microliters of sample were injected, and the flow rate was set at 0.4 mL/min. Electrospray ionization (ESI) was selected as the ionization source for QTOF-MS analysis (30). Raw data from UHPLC-QTOF-MS were analyzed using analysis software. First, features were extracted from the raw data using Agilent MassHunter Profinder software 10.0. Normalization and library-based metabolite annotations were performed using Mass Profiler Professional software and MassHunter METLIN Metabolite Personal Compound Databases and Libraries. Annotated metabolome data were then statistically analyzed using MetaboAnalyst 6.0 and finally visualized using R (version 4.3.1).

### RNA-sequencing

For transcriptomic experiments, a commercially available reconstructed human pigmented epidermis model (SkinEthic™ RHPE, Phototype VI, ITA < −40°; EpiSkin, France) was used. The 0.5-cm² model consisted of normal human keratinocytes and melanocytes derived from foreskin (batch 24-RHPE-030_S) (31). The models were stabilized by incubating at 37°C in a 5% CO₂ environment for 24 hours. Following treatment, total RNA was extracted from four RHPE groups: (1) the non-treated control, (2) 100 nM of α-MSH, (3) 100 nM of α-MSH with 100 nM of α-MSH + DMSO, and (4) 100 nM of α-MSH with 4 µM of ellagic acid. RNA quantity and quality were assessed before library preparation. Messenger RNA was enriched from 2 μg of total RNA using oligo(dT) magnetic beads, followed by fragmentation and complementary DNA synthesis. Sequencing libraries were prepared using an Illumina-compatible stranded mRNA library preparation kit according to the manufacturer’s instructions. The library preparation procedure included end repair, 3′ adenylation, Illumina adapter ligation, library amplification, and size selection.

The quality and fragment-size distribution of the final libraries were evaluated using an Agilent 2100 Bioanalyzer or TapeStation system. Qualified libraries were sequenced on an Illumina sequencing platform to generate 150-bp paired-end reads.

### Analysis of differentially expressed genes (DEGs)

Raw sequencing data were assessed using FastQC (https:// www.bioinformatics.babraham.ac.uk) and subjected to a quality control step to discard low-quality reads via Trimmomatic (32). This step included discarding reads with more than 10% skipped bases (marked as ‘N’s), sequencing reads with over 40% of bases possessing a quality score less than 20, and those with an average quality score below 20. Only high-quality reads were mapped to the human reference genome (Homo sapiens: GRCh38) using the Spliced Transcripts Alignment to a Reference (STAR) aligner (33). For downstream DEG analysis, only uniquely mapped read pairs were utilized. Gene expression levels were quantified by RNA-Seq by Expectation-Maximization (RSEM v1.3.3). Differential expressions were analyzed using DESeq2 in R. DEGs with a log2 fold-change (log2 FC) greater than one and an adjusted p-value (q-value) less than 0.05 were considered statistically significant. The overall expression pattern was visualized through pairwise correlation analysis, scatterplots, hierarchical sample clustering heatmaps, and principal component analysis plots using the ggplot2 R package. Heatmap clustering analysis of DEGs was performed based on log2 FPKM values using hclust2 package (v3.6.2; available at https://github.com/SegataLab/hclust2), employing Pearson’s distance for clustering distance calculation and complete linkage method for hierarchical clustering.

### Functional enrichment and interaction network analysis

K-means clustering was used to group genes based on their expression patterns altered by extrinsic treatment. Gene Ontology (GO) terms and enrichment analyses of DEGs were performed using Metascape (http://metascape.org/gp/index.html). The Metascape analysis workflow adhered to the following criteria: Initially, multiple gene lists identified from the DEG analysis were used as input. For GO annotation, three primary categories of gene functions - cellular component (CC), molecular function (MF), and biological process (BP) - were extracted. Subsequently, functional enrichment analysis was conducted using default parameters: a minimum overlap of three, an enrichment factor of 1.5, and a p-value threshold of 0.01 for filtering purposes (34).

Protein-protein interactions (PPIs) related to skin barrier molecules were mapped using the Search Tool for Retrieval of Interacting Genes (STRING; https://string-db.org), retaining only nodes that had high confidence interaction scores (> 0.7).

## Acknowledgements

The authors of the research paper acknowledged the Center for Bio-Medical Engineering Core Facility at Dankook University for providing valuable equipment, including a LightCycler 480 II quantitative real-time PCR (NFEC-2015-10-205658), and for providing space.

## Author Contributions

S.M.K., H.W.J, and S.M. contributed to the conceptualization and methodology of the study. S.M.K. and H.W.J performed the experimental investigations, including bacterial isolation and characterization, cell-based efficacy assays, and validation experiments. S.M. and H.W.J contributed to experimental design, formal data analysis, transcriptomic and bioinformatic analyses, data interpretation, and visualization. M.K. and M.-J.K. contributed to bacterial culture, cell-based experiments, and experimental validation. D.-G.L. contributed to methodology, metabolomic analysis, data interpretation, and validation. S.K. contributed to study design, experimental methodology, data analysis, and interpretation. H.W.J. and K.H. contributed to conceptualization, supervision, project administration, and interpretation of the results. S.M.K. and S.M. wrote the original draft of the manuscript. S.M., D.-G.L., S.K., H.W.J., and K.H. reviewed and edited the manuscript. H.W.J. and K.H. jointly supervised the study. All authors reviewed and approved the final version of the manuscript.

## Ethics declarations

### Ethics approval and consent to participate

The collection of facial skin samples from the healthy participant for microbiota isolation was approved by the Institutional Review Board of H&BIO Corporation R&D Center (IRB Protocol No. HBABN01-210217-HR-0181-01), and the study was conducted in accordance with the principles of the Declaration of Helsinki.

For the transcriptomic experiments, a commercially available reconstructed human pigmented epidermis model (SkinEthic™ RHPE, Phototype VI; EpiSkin, France) was used. The model consists of normal human keratinocytes and melanocytes derived from foreskin and was provided as a commercially manufactured human-derived tissue model for in vitro research. No identifiable donor information was obtained or accessed by the investigators.

### Data availability statement (mandatory)

The datasets generated and analyzed during the current study, including the data supporting the experimental and transcriptomic findings, are available from the corresponding author upon reasonable request.

## Additional Information (including a Competing Interests Statement)

### Consent for publication

Not applicable.

